# SpermQ - a simple analysis software to comprehensively study flagellar beating and sperm steering

**DOI:** 10.1101/449173

**Authors:** Jan N. Hansen, Sebastian Raßmann, Jan F. Jikeli, Dagmar Wachten

**Affiliations:** Institute of Innate Immunity, Biophysical Imaging, University Hospital Bonn, University of Bonn, 53127 Bonn, Germany; Center of Advanced European Studies and Research (caesar), Molecular Physiology, 53175 Bonn, Germany

## Abstract

Motile cilia, also called flagella, drive cell motility across a broad range of species; some cilia propel prokaryotes and eukaryotic cells like sperm, while cilia on epithelial surfaces create complex fluid patterns e.g. in the brain or lung. For sperm, the picture has emerged that the motile cilium, also called flagellum, is not only a motor, but also a sensor that detects stimuli from the environment, computing the beat pattern according to the sensory input. Thereby, the flagellum of the sperm cell navigates the sperm through a complex environment like the female genital tract. However, we know very little about how environmental signals change the flagellar beat and, thereby, the swimming behaviour of sperm. It has been proposed that distinct signalling domains in the flagellum control the flagellar beat. A detailed analysis has been mainly hampered by the fact that current comprehensive analysis approaches rely on complex microscopy and analysis systems. Thus, knowledge on sperm signalling regulating the flagellar beat is based on custom quantification approaches that are limited to only a few aspects of the flagellar beat, do not resolve the kinetics of the entire flagellum, rely on manual, qualitative descriptions, and are little comparable among each other. Here, we present *SpermQ*, a ready-to-use and comprehensive analysis software to quantify sperm motility. *SpermQ* provides a detailed quantification of the flagellar beat based on common time-lapse images acquired by dark-field or epi-fluorescence microscopy, making *SpermQ* widely applicable. We envision *SpermQ* becoming a standard tool in flagellar and motile cilia research that allows to readily link studies on individual signalling components in sperm and distinct flagellar beat patterns.

## Introduction

Reproduction starts with fertilization, the fusion of sperm and egg. In mammals or other internal fertilizers, sperm need to pass a complex environment, the female genital tract, to reach the site of fertilization. On their way through the female genital tract, sperm cells are guided by different physical and chemical cues. Here, the flagellum, the sperm tail, functions as both, sensor and motor of the sperm cell, allowing guided motility to the site of fertilization (Wachten et al., 2017). Several physical and chemical cues guide the sperm cell through the complex environment in the female genital tract. Chemotaxis, thermotaxis, and rheotaxis have been described to control the sperm’s swimming path (Wachten et al., 2017), whereby the sperm flagellum serves as a sensor and motor, transducing the sensory input from the environment into a distinct flagellar beat pattern (Alvarez et al., 2014).

A symmetrical flagellar beat results in a straight swimming path of the sperm cell, whereas asymmetries in the beat pattern lead to a curved or even helical swimming path (Friedrich et al., 2010; Goldstein, 1977; Jikeli et al., 2015; Rikmenspoel, 1965; Rikmenspoel et al., 1960). Thus, control of beat asymmetry in sperm might be the key regulator of steering and navigation. In fact, in sea urchin sperm, the symmetry of the flagellar beat pattern determines the swimming path of the sperm cell (Jikeli et al., 2015). However, the navigation of mammalian sperm is less well understood. In bull sperm, an asymmetric flagellar beat pattern has been directly correlated to a curved or rotating swimming path (Rikmenspoel et al. 1960; Rikmenspoel 1965; Friedrich et al. 2010). In human sperm, analyzing the beat pattern in 3D revealed that during rheotaxis, sperm swimming on a curved trajectory display an asymmetric beat in the midpiece (Bukatin et al., 2015). Many studies unravelled only selected aspects of the flagellar beat, investigating only a small part of the flagellum (Jansen et al., 2015; Krähling et al., 2013), or analysing the swimming path and swimming velocity only (Chen et al., 2016; Hirano et al., 2003; Holt et al., 1994; Wang et al., 2003). In addition, the current perspective on sperm signalling points towards a compartmentalization of signalling along the flagellum, e.g. in the case of cAMP signalling (Balbach et al., 2017; Mukherjee et al., 2016).

Thus, a detailed analysis of the flagellar beat is needed to understand sperm navigation. To this end, we have developed *SpermQ*, an analysis software that allows to comprehensively study the flagellar beat pattern and sperm steering using microscopy techniques that can be readily applied by almost any lab, i.e. dark-field and epifluorescence microscopy. *SpermQ* cannot only analyse the beat of tethered sperm, but also the beat of free-swimming sperm. We envision *SpermQ* becoming a standard tool in sperm research.

## Results

### Analysis and workflow performed by the software *SpermQ*

*SpermQ* is an automated analysis software to comprehensively study the flagellar beat of sperm in time-lapse image sequences generated by dark-field microscopy. Initially, *SpermQ* determines the flagellar trace in every frame of the analysed image sequence. First, the raw image (Figure 1A) is gauss-filtered, segmented, and skeletonized to obtain a rough trace of the flagellum (Figure 1B). Next, along the detected flagellar trace, normal lines are constructed (Figure 1C). The image intensities along each normal line are fitted to a Gaussian curve. The detected flagellar trace is adjusted to the centre of the Gaussian curve, as this defines the precise location of the flagellum in the image (Figure 1D).

**Figure 1:**
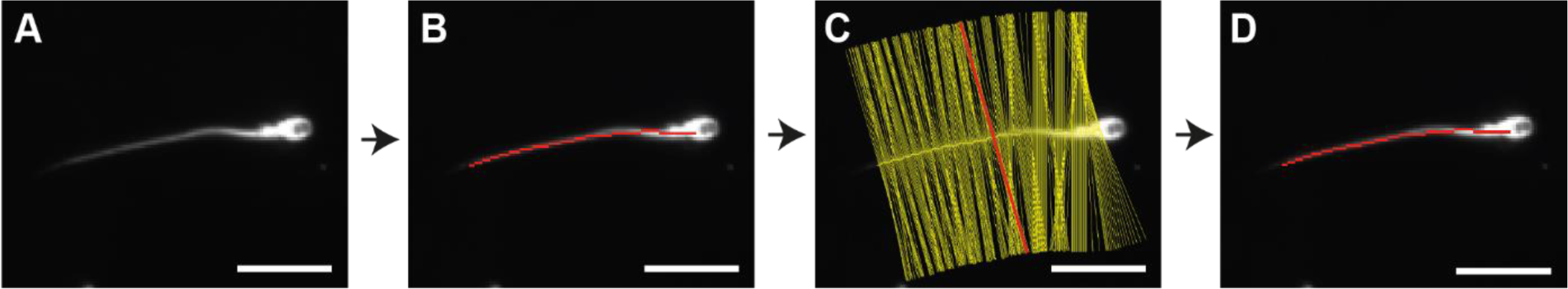
Reconstruction of the flagellum. **(A)** Each frame image is segmented and skeletonized to obtain **(B)** a rough track of the flagellum. **(C)** Along the flagellum, tangential vectors are determined to generate normal lines (yellow; red: exemplary normal line). The intensity profile along each normal line is fitted to a Gaussian curve to obtain the centre of the profile. The track points are then corrected by shifting them to the centre, **(D)** resulting in a precise flagellar track. Bar: 20 μm.

After reconstructing the flagellum, *SpermQ* determines **orientation parameters** (Figure 2A-C) and **flagellar parameters** (Figure 2D-H).

**Figure 2:**
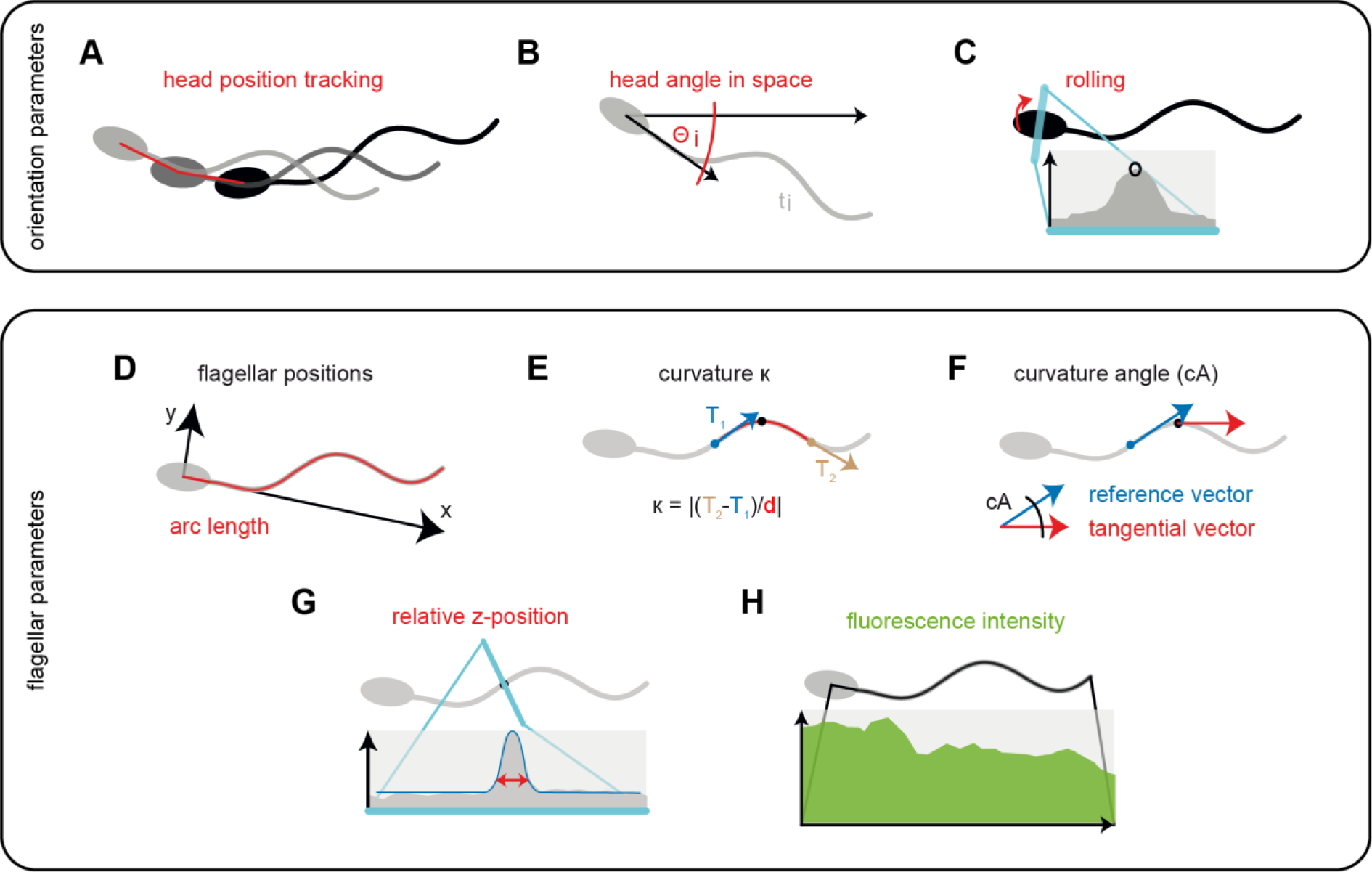
*SpermQ* output parameters. *SpermQ* outputs parameters characterize the orientation of the whole cell in space (A-C) or each individual position on the flagellum (D-H). *SpermQ* outputs the head position in space over time (A) and the angle Θ (B), which is the angle of the head-midpiece-axis to the x-axis. The maximum intensity (black circle) within the first normal lines of the flagellar track describe sperm rolling (C). To determine x- and y-positions of the flagellar points relative to the head, all flagellar points are translated into a coordinate system originating from the center of the head (D): the initial 6 pm of the flagellum define the x-vector of the coordinate system. The curvature of a given point on the flagellum is determined as the geometric curvature and by using the tangential vectors 5 μm before and after the point (E). For arc lengths close to the head or the flagellar tip, a smaller distance is chosen. The curvature angle (cA) at a given point is calculated by the angle between the tangential vector at the given point and the tangential vector of the point 10 μm before the given point (F). As a relative parameter for the flagellar z-position, *SpermQ* outputs the width of the Gaussian curve fitted to the intensity profile of the normal vectors at every point of the flagellum (G). Furthermore, the maximum intensity of every flagellar point is measured to generate an intensity profile along the flagellum (H).

The **orientation parameters** describe the location and orientation of the sperm cell in its environment and allow studying sperm navigation. Orientation parameters are determined using the initial points of the reconstructed flagellar trace. *SpermQ* defines the head of the sperm cell, representing the first point of the detected flagellar trace, as its position in space (Figure 2A). This allows to track the swimming path of the sperm cell over time and thus, to determine whether the sperm cell swims straight or on a curved path. The head angle in space (Θ, Figure 2B) reveals the orientation of the sperm cell. Θ is defined as the angle between the head-midpiece-vector and the x-axis of the image. The oscillation of Θ determines the main beat frequency. The maximum intensity in the head serves as a parameter to determine rolling of the sperm cell around its longitudinal axis (Figure 2C). Due to the elliptical and slightly asymmetrical geometry of the sperm head, the intensity of the light that is scattered from the head into the direction of the objective during dark-field imaging oscillates while the sperm cell is rolling. Determining the orientation parameters over time allows to reveal the trajectory of the swimming path, the curvature of the trajectory, the rolling of the sperm cell around its own axis, and the main beat frequency of the sperm.

The **flagellar parameters** link the flagellar beat pattern to the swimming path and describe the location, intensity, and local curvature of individual points on the flagellum. A position on the flagellum is defined by its arc length, which is the distance of the given point to the head on the stretched flagellum. In each frame, all points along the flagellum are translated into a new coordinate system, whose x-axis is defined as the vector through the head and the midpiece of the sperm (Figure 2D). Calculating the range of the x- and y-coordinates of the flagellum relative to the head over time reveals the flagellar envelope, which refers to the beat amplitude. Additionally, the symmetry of the flagellar envelope compared to the head axis is important as this controls the swimming path of the sperm cell.

To determine the location and propagation of waves along the flagellum, two different parameters were included, which describe the local curvature of the flagellum, independently of the head axis: the curvature (Figure 2E) and the curvature angle (Figure 2F). The curvature at a given arc length is determined based on the equation for the geometric curvature and using tangential vectors at defined distances (e.g. 5 μm) before and after the given arc length. The curvature angle at a given arc length is defined as the angle between the tangential vector at and slightly before (e.g. 10 μm) the given arc length.

To consider that the flagellar beat shows also a component that is vertical to the focal plane of the microscope, we implemented a method that qualitatively describes the position of the flagellum along the optical axis of the microscope, representing the z-axis. The focal position *o* along the optical axis in the specimen that is recorded by the microscopy set-up is, according to the lens equation, defined as *o* = (*f* · *i*)/(*i* − *f*), where *i* is the position of the conjugate image generated by a lens with a focal length *f*. In a microscopy system, *f* is mainly represented by the objective, determining the focal position in the specimen. When an object is located at the focal position, it appears sharp in the image that is recorded by the microscopy set-up. The more distant to the focal position along the optical axis the object is, the less sharp and the wider it appears in the recorded image. This approach has been proposed already in the 19^th^ century to qualitatively describe the position of the flagellum on the optical axis, representing the z-position (Rikmenspoel et al., 1960). Recently, this method was applied to describe the relative z-position of the flagellum at a given point by the width of the intensity peak on the normal line of the given flagellar point (Bukatin et al., 2015). Similarly, we included an algorithm into *SpermQ* that fits the intensity profile of every normal line along the flagellum to a Gaussian curve, whose width parameter represents the z-position of the object (Figure 2G). The relative z-position, determined by *SpermQ* is not calibrated and does not reveal whether the object is located above or below the focal plane and, thus, the respective results have to be interpreted carefully. A way to solve this, however, has been published elsewhere (Bukatin et al., 2015), but has not been implemented in *SpermQ.*

To analyse fluorescently labelled sperm, *SpermQ* determines the fluorescence intensity at every arc length and in every frame as the maximum of the Gaussian curve fitted to the normal intensity profile (Figure 2H). This allows localizing fluorescent molecules in the flagellum and determining the fluorescence intensity along the flagellum. Of note, also the z position of the flagellum influences intensity measurements. However, this influence can be neglected when applying bright fluorescent labels or when filtering out intensity oscillations in the range of the flagellar beat frequency.

For every flagellar parameter, *SpermQ* generates a kymograph, which displays the parameter results intensity-coded as a function of arc length and time, depicting the whole kinetics of an individual parameter at every flagellar position (e.g. Figure 3D).

**Figure 3:**
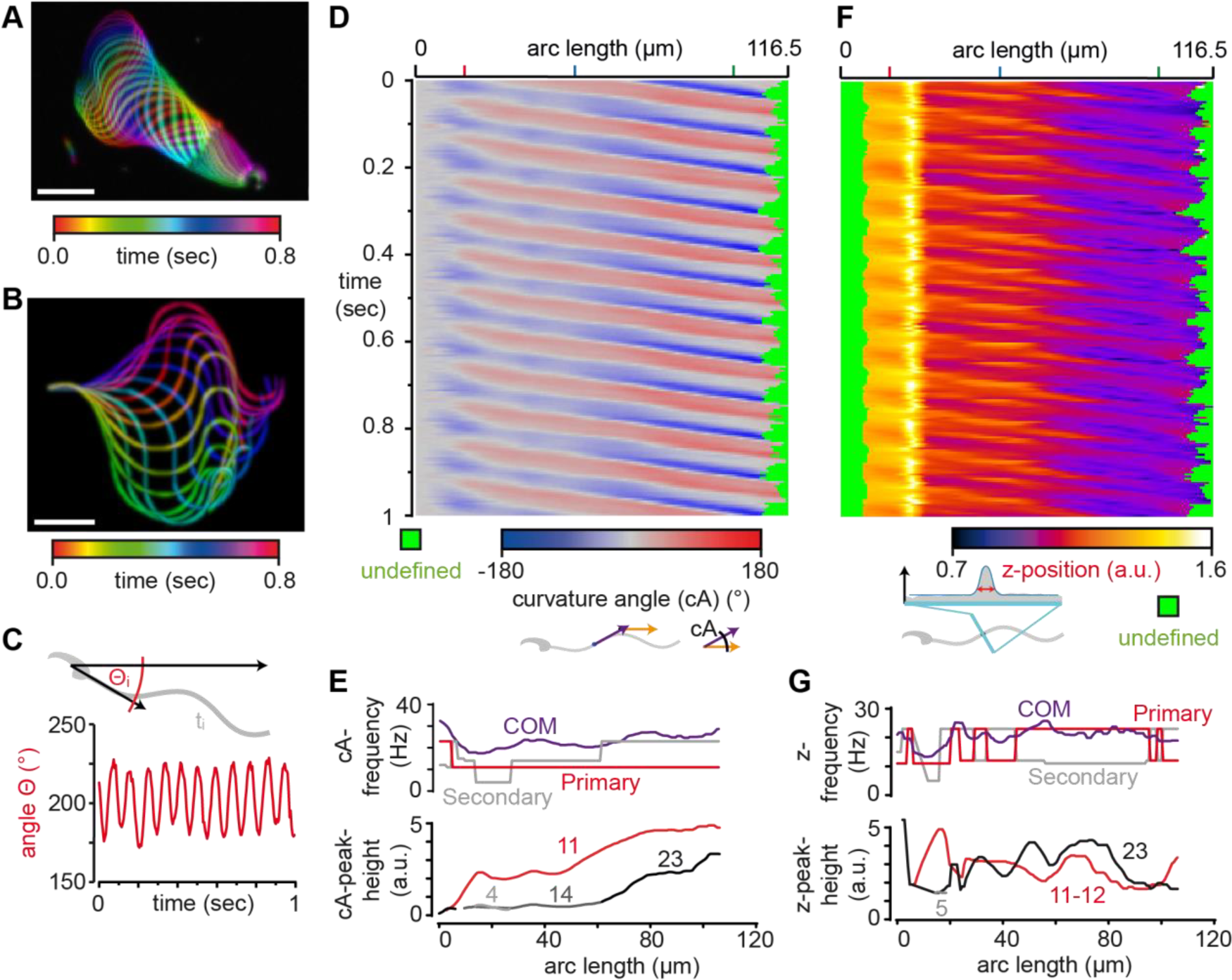
*SpermQ* reveals the beat pattern of a head-tethered mouse sperm. *SpermQ* analysis of a wild-type mouse sperm, tethered to a glass surface at the head. Projection of the raw image (A) and the trace after orientation to the head-midpiece-axis (B), each representing one beat cycle. *SpermQ* analysis reveals that the head-orientation angle Θ stably oscillates with a primary frequency of 11 Hz around an angle of 200° in space (center of mass (COM) of frequency spectrum: 19.9 Hz) (C). Kymographs of the determined curvature angle (cA) (D) or the z-position relative to the plane (F). *SpermQ* frequency analysis of the cA (E) or of the z-position (G) along the flagellum. Amplitudes for selected peaks in the frequency spectrum (frequency values indicated in Hz) are plotted in the bottom graph. Scale bars: 20 μm.

The flagellar beat of the sperm cell is highly periodic and thus, can be considered as an oscillation. To determine the beat frequency of sperm, Fast Fourier Transformation (FFT) of the oscillating parameter and analysis of the resulting frequency spectrum is commonly used to determine the beat frequency and other beat characteristics, such as the phase of the oscillation or additional frequencies on the flagellum that control the swimming path (Saggiorato et al., 2017). To implement this method into *SpermQ*, we included a FFT algorithm (derived from the package *edu.emory.mathcs.jtransforms.fft* by Piotr Wendykier, Emory University). The FFT algorithm is used to translate the time course of each parameter– and for the flagellar parameters also at each flagellar position – into a frequency spectrum. For each frequency spectrum, *SpermQ* determines the positions of the highest (primary frequency) and the second highest (secondary frequency) peak in the spectrum. In addition, *SpermQ* determines the center-of-mass (COM) of the frequency spectrum. The COM describes the distribution and amplitude of all frequency peaks in the frequency spectrum. As the COM depends on the characteristics of the whole frequency spectrum, the COM serves as a quick and simple measure to indicate differences in the frequency spectra of multiple cells (or over time). Large difference in the COM highlight that the user should analyse the respective frequency spectra in more detail.

In summary, *SpermQ* determines the location and orientation of the cell in space, the precise beat pattern along the flagellum, the intensity levels on the flagellum, and the frequency characteristics of all of these parameters. An overview of all parameters is presented in Table 1.

**Table 1.** Terminology and indications of *SpermQ* parameters

| Parameter | Indication |
| --- | --- |
| **Arc length** | position on the flagellum |
| **head position tracking** | trajectory of the swimming path |
| **head angle in space ($\Theta$)** | main beat frequency, orientation in space, curvature of the trajectory |
| **maximum intensity in head (head torsion)** | main beat frequency, rolling frequency (free-swimming sperm) |
| **x coordinate** | oscillations on the longitudinal axis through the head |
| **y coordinate** | beat amplitude, asymmetry of the whole beat waveform |
| **Curvature / curvature angle** | localization of the origin and propagation of waves on the flagellum, wave kinetics, beat frequency, local beat amplitude, local beat asymmetry |
| **z position** | qualitative value to describe the kinetics of the flagellar beat in the direction vertical to the focal plane |
| **intensity** | intensity profile of the cell along the whole flagellum, instructive when the cell is labelled with a fluorescence indicator; intensity also depends on the position of the flagellum in z |

### *SpermQ* analysis of the beat of a head-tethered mouse sperm

To demonstrate the power of *SpermQ*, we analysed dark-field microscopy images of a head-tethered mouse sperm (Figure 3). The orientation of the sperm cell to the head-midpiece-axis (Figure 3B) demonstrates that the flagellar envelope is symmetric. Analysis of Θ reveals that the cell beats regularly with a beating frequency of 11 Hz (Figure 3C) without rotating around the tethered point (the overall Θ level is stable over time). The homogeneity of the beat pattern is confirmed by the curvature angle kymograph (Figure 3D). Based on the kymograph, *SpermQ* detects a primary frequency peak of 11 Hz along the entire flagellum and a secondary frequency peak of 23 Hz close to the flagellar tip (Figure 3E, bottom panel). This shows that *SpermQ* is able to resolve the flagellar beat pattern spatially along the flagellum. The same flagellar beat frequencies are apparent in the kymograph that depicts the relative z position of the flagellum (Figure 3F). Of note, this kymograph displays multiple maxima/minima for one oscillation at the tip of the flagellum (Figure 3F). This is due to the lack of direction information compared to the focal plane in the parameter relative z position. When the flagellum crosses the focal plane during its beat cycle, a single beat will result in a double frequency peak. As a result, in the frequency analysis, the double frequency of the primary beat frequency is detected and adds on the power of the second harmonic frequency in the spectrum, resulting in the false detection of the second harmonic frequency as the primary frequency peak (Figure 3G). This example highlights that the relative z position needs to be carefully interpreted. However, this shortcoming can be avoided by setting the imaging focal plane within the cover glass adjacent to the tethered cell that is recorded.

### *SpermQ* analysis of the beat of a freely-swimming human sperm

*SpermQ* is also able to analyse the beat and swimming path of freely-swimming sperm, e.g. in dark-field microscopy images of human sperm (Figure 4). Head-tethered sperm are not able to roll around their longitudinal axis and thus, their flagellar beat is mainly parallel to the focal plane. In contrast, freely-swimming sperm role around their longitudinal axis while beating in all different directions. The 2D projection of one beat cycle of a freely-swimming human sperm cell displays an asymmetric shape of the beat (Figure 4A), even when aligning the sperm cell to the head-midpiece axis in 2D (Figure 4B). Here, the rolling frequency is 9 Hz and the Θ results show that two periodic events determine the flagellar beat frequency measured in 2D (Figure 4B): a primary frequency of 9 Hz (which corresponds to the rolling frequency) and a secondary frequency of 27 Hz (= three times the primary frequency). This indicates that while the sperm is rolling once around its longitudinal axis, the flagellum performs three beat cycles. By tracking the head position, *SpermQ* creates trajectories of the swimming path (Figure 4D) and determines a precise kymograph of the curvature angle along a freely-swimming sperm (Figure 4E). Taken together, *SpermQ* precisely quantifies the beat of freely-swimming sperm.

**Figure 4:**
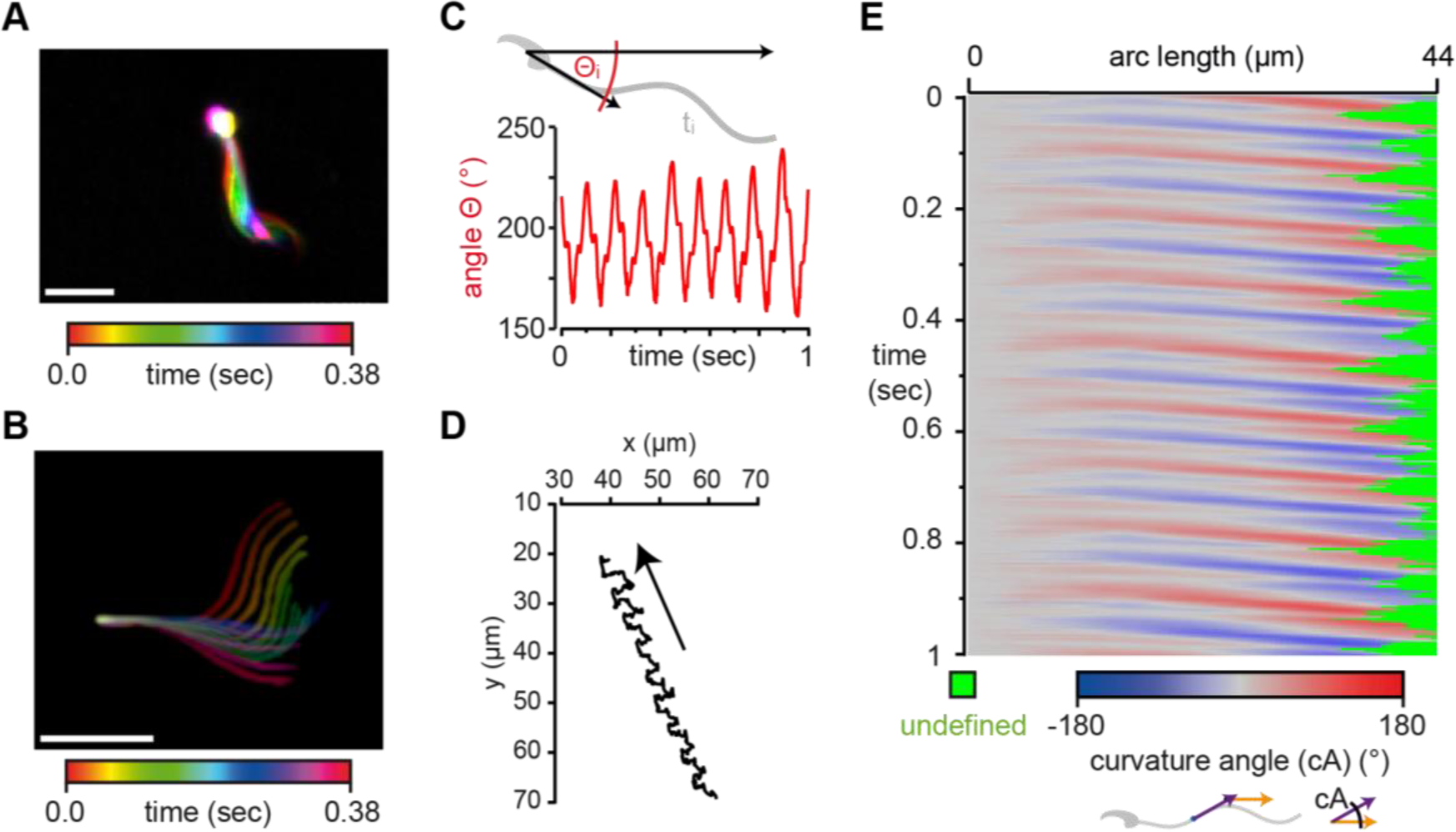
*SpermQ* reveals the beat pattern of a free-swimming human sperm. *SpermQ* analysis of a human sperm, freely-swmming in a shallow observation chamber (depth 300 μm). Projection of the raw image (A) and the trace after orientation to the head-midpiece-axis (B), each representing one beat cycle. SpermQ-analysis reveals that the head-orientation angle Θ stably oscillates with the primary frequency of 9 Hz (rolling frequency of the sperm) and the secondary frequency of 27 Hz (beat frequency) (center of mass (COM) of frequency spectrum: 21.2 Hz) (C). Trajectory of the sperm’s head position. Kymograph of the determined curvature angle (cA) (E). Scale bars: 20 μm.

### *SpermQ* is a readily-applicable software

We developed *SpermQ* with the goal to create a software that is easy to handle, fast, allows batch processing, and requires as little user interaction as possible. *SpermQ* is a completely automated software.

Before applying the software, it is recommended to split the image stack to-be-analyzed into single-sperm image stacks to exclude other particles (e.g. dirt particles, other sperm) from the analyzed image. This also increases the analysis speed by reducing the data size of the processed image. Each of the single-sperm image stacks needs to be cropped to the minimum required size, but leaving a little bit of extra space (few microns) around the flagellum. The extra space is required for Gaussian curve fits along the flagellum (Figure 1C). Before loading these stacks into *SpermQ*, the background in the image has to be reduced. To this end, the “Subtract Background” method in ImageJ can be applied, which, when a low radius is selected in the “Subtract Background”-settings, also equalizes intensity differences in the sperm cell, thereby improving flagellar reconstruction by *SpermQ*.

After preprocessing, the data can be readily applied to *SpermQ*. When launching *SpermQ*, a settings dialogue opens to set default settings for distinct types of data sets (e.g. depending on the sperm species or the magnification used). To load data into *SpermQ*, a dialogue containing an analysis list opens, to which the user can add single-sperm image stacks from the hard disk. After listing all sperm image stacks, the software can be started. In the first step, time projections of each selected image stack are presented, in which the user is asked to set a ROI. This ROI should contain an image region where the sperm cell is always present in the entire sequence. For tethered sperm, a circle around the head is sufficient. For freely-swimming sperm, a ROI around the sperm’s swimming path is set. Setting this ROI reduces the regions where *SpermQ* searches for a flagellum in the image and, thereby, increases analysis speed and precision of the software, as e.g. remaining dirt particles in the image will be neglected. After setting the ROIs, *SpermQ* analyzes all image stacks without further user interaction. The results are automatically saved into a newly created analysis folder, separately for each analyzed sperm.

To visualize the results, we developed an additional open-source java application, the *SpermQ Evaluator* (https://github.com/IIIImaging/SpermQ Evaluator). In *SpermQ Evaluator*, data sets can be loaded and ordered by the user, and overview tables for the loaded data sets are created. In addition, *SpermQ Evaluator* saves an overview PDF for each analyzed data set, comprising a collection of plots of the most important parameters (see Supplementary File 1).

## Discussion

How mammalian sperm control their flagellar beat pattern and thereby, their swimming path still remains elusive. The detailed analysis of the flagellar beat pattern during sperm navigation has been hampered by the lack suitable software and hardware solutions that are widely applicable. A plethora of custom-made imaging and image analysis approaches have been used that were limited in the number of parameters or the fraction of the flagellum that could be analysed, or that were restricted to sperm velocity parameters only (e.g. CASA).

With *SpermQ*, we developed a software that analyses the flagellar beat in detail while requiring only simple imaging approaches (dark-field or epifluorescence microscopy). *SpermQ* analyses the flagellar beat in depth, unravelling different orientation parameters and flagellar parameters, including a FFT-based frequency analysis of all parameters. In combination with the evaluation tool *(SpermQ Evaluator)*, comprehensive results can be rapidly and clearly displayed. Furthermore, *SpermQ* is automatized, allowing to analyse large data-sets and excluding the bias of user-dependent custom analysis approaches. By providing default settings, *SpermQ* is readily applicable by any user. We even included the tracking of not only tethered sperm but also free-swimming sperm in *SpermQ*. Thereby, we make it easily applicable to study flagellar beating in free-swimming sperm, even for labs that did not have the know-how and technical requirements to perform such complex tracking so far.

Taken together, we envision that *SpermQ* makes studies on the flagellar beat faster, ready to perform by any user, and more comparable.

## Material and Methods

### Mouse sperm

Wild-type mice (C57BL/6) were obtained from Janvier Labs (France). Animal care and experiments were in accordance with the relevant guidelines and regulations and approved by the local authorities (LANUV). Mice were killed by cervical dislocation after isoflurane (Curamed Pharma) inhalation. Mouse sperm were isolated by incision of the cauda epididymis followed by a swim-out in modified TYH medium (in mM: 135 NaCl, 4.8 KCl, 2 CaCl2, 1.2 KH2PO4, 1 MgSO4, 5.6 glucose, 0.5 sodium pyruvate, 10 lactic acid, 10 HEPES, pH 7.4 adjusted at 37°C with NaOH). After 15 - 30 min swim-out at 37°C, sperm were collected and counted.

### Human sperm

Human semen samples were donated by healthy adult males with their prior written consent and the approval of the ethic committee of the University of Bonn (042/17). The sperm cells were purified by a “swim up” procedure (Strunker et al., 2011) using human tubular fluid (HTF) (in mM: 97.8 NaCl, 4.69 KCl, 0.2 MgSO_4_, 0.37 KH_2_PO_4_, 2.04 CaCl_2_, 0.33 Na-pyruvate, 21.4 lactic acid, 2.78 glucose, 21 HEPES, and 25 NaHCO3 adjusted to pH 7.3 - 7.4 with NaOH.

### Imaging

Dark-field imaging was performed at an inverted microscope (IX71; Olympus) equipped with a dark-field condenser and a high-speed camera (PCO Dimax). Image sequences of human sperm were recorded with 500 frames per second (fps) and using a 20x objective (NA 0.5, UPLFLN; Olympus); image sequences of mouse sperm were recorded with 200 fps and using a 10x objective (NA 0.4, UPlanFL; Olympus) with an additional 1.6x magnifying lens (Olympus) that was inserted into the light path (final magnification: 16x). For imaging, the sperm solution prepared by the swim-up technology was inserted in a custom-made observation chamber with a depth of about 150 μm. The temperature of the microscope incubator (Life Imaging Services, Basel, Switzerland) was adjusted to 37°C for imaging.

### Image analysis

All image processing and analysis was performed in ImageJ (U.S. National Institutes of Health, Bethesda, Maryland, USA). For images of mouse sperm, a minimum-intensity projection of each image sequence was generated and subtracted from the respective image sequence. Thereby, background signals were reduced in the image. In single-plane images of human sperm, the background was removed by the ImageJ function *Subtract Background* (radius: 10 px for a magnification of 20x). After background correction, image sequences of mouse and human sperm were subjected to *SpermQ* analysis (settings: Table 2).

**Table 2.**
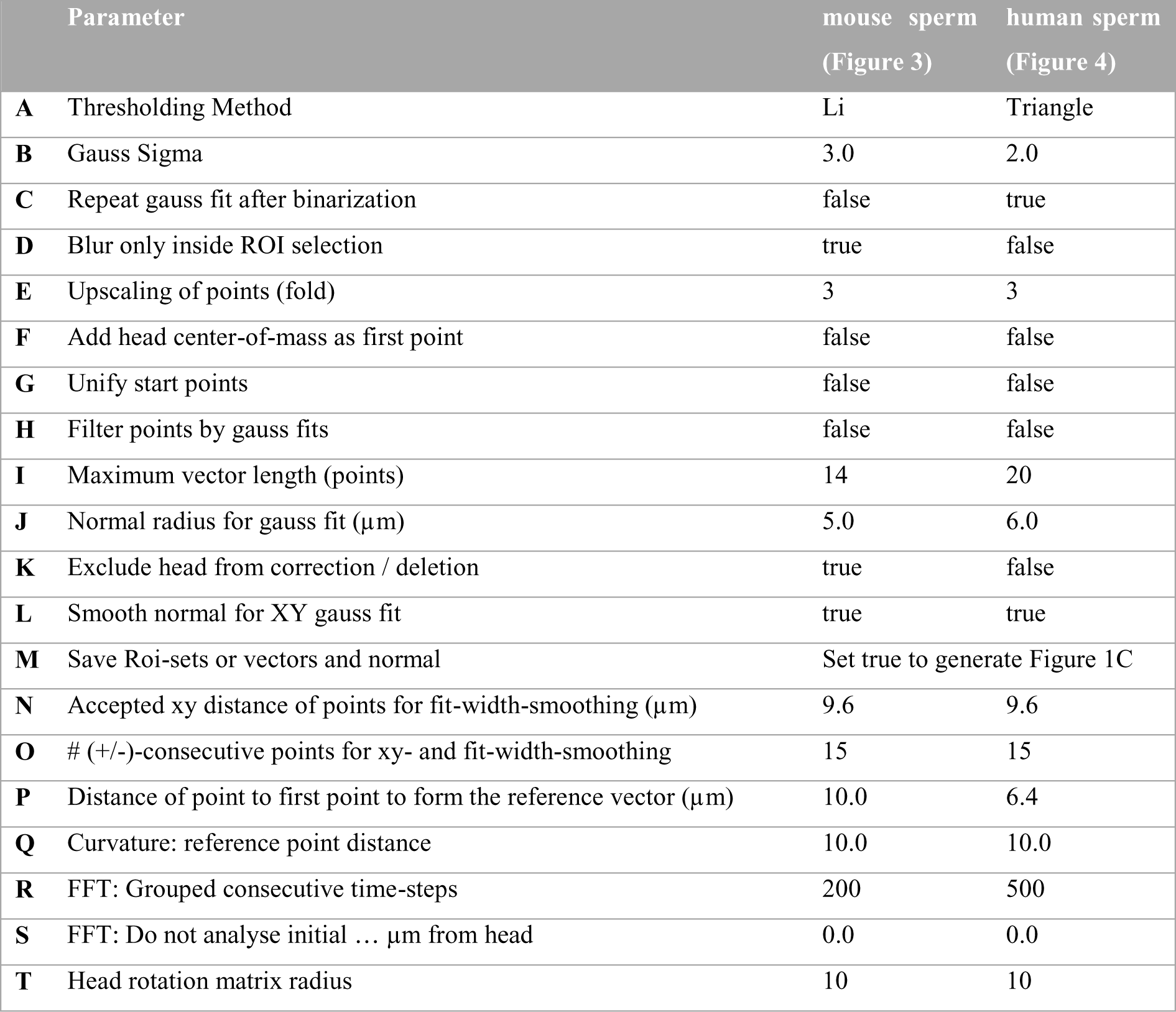
*SpermQ* settings

### Analysis workflow underpinning *SpermQ*

*SpermQ* is a fully automated analysis software to quantify sperm navigation and the flagellar beat of sperm. *SpermQ* was developed as an ImageJ plugin in Java. We will present the workflow performed by *SpermQ* in the following. The parameters applied in the *SpermQ* workflow are listed in Table 2 and will be referenced at the steps where they determine the workflow. For each image sequence, each time frame is processed separately. To reconstruct the flagellum, the frame image is filtered by a Gaussian kernel with a σ defined by the user (Table 2B), binarized using one of the thresholding algorithms in ImageJ selected by the user (Table 2A), again Gaussian blur filtered with the same σ (Table 2B-C), and processed by the ImageJ plugin *Skeletonize3D* (Arganda-Carreras et al., 2010). The Gaussian blur can be restricted to the ROI set by the user, e.g. to blur only the sperm head while preserving fine structure of the flagellum (Table 2D). The generated skeleton image is analysed (*AnalyzeSkeleton;* Arganda-Carreras et al. (2010)) to retrieve the pixels of the skeleton. For each pixel, a point *p_i_* is generated using the coordinates of the pixel and all points are collected in a point list *P* (the index *i* indicates the position of the point in *P*; 0 < *i* ≤ |*P*|). Next, *P* is sorted so that 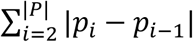 is minimal. To ensure that *p*_1_ corresponds to the sperm head, we take advantage of the fact that dark-field images are brighter at the head due to a stronger light scattering than at the tip of the flagellum. The average image intensities in circular ROIs (radius: 8 pixels) around *p*_1_ and around *p*_|*P*|_ are compared and the order of *P* is inverted if the average intensity around *p*_1_ is smaller than around *P*_|*P*|_. SpermQ can be set to add the center of mass of the circular ROI as the first point of *P* (Table 2F) or to always start at the same point as the first point in each frame (Table 2G). For the latter, *p*_1_ is set in each frame to the average position of all *p*_1_ in all frames across the time series.

Next, to obtain a finer flagellar curve, |*P*| is increased by linear interpolation between its elements: between two consecutive points (*p_i_* and *P_i+1_*), a distinct number of equidistant points is inserted. The number of points is set by the user (Table 2E); by default 2 points are inserted between each pair of *p_i_* and *p_i+1_*. Next, for each point *p_i_*, the tangential vector 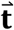 pointing towards *p_i+1_* is determined. The tangential vector 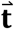 is determined by the vector between the points located a given number of points (Table 2I, divided by two) before and after *p_i_*. At the start or the end of *P*, closer points are selected to form 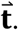. The normal vector 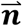 was determined as 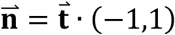. The sign convention used for normal direction was such that if the cross product 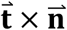 was positive, 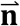 was multiplied by −1. The normal vector 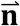 was used to generate a normal line with a defined length (Table 2J) from *p_i_* towards both orthogonal directions compared to 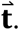. Next, the intensity profile along the normal line was determined in steps of a pixel length (for a 20x magnification and an 11 μm camera pixel size: 0.55 μm / pixel). Each intensity value *i_n_* along the normal line is linearly interpolated from the four surrounding pixels weighted by their distance to the point on the normal line. The intensity profile is smoothed, if set so (Table 2J), and then fitted to a Gaussian curve using the *CurveFitter* implemented in ImageJ (maximum number of iterations: 1000). If the determined parameters of the Gaussian curve fulfilled all following four criteria, the point *p_i_* was shifted by 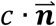 (*c* = centre of the fitted Gaussian curve) to center *p_i_* on the flagellum:

1. *r*^2^ > 0.8 (*r*^2^ = quality of the fit)
2. *a* > 0 (*a* = height the Gaussian curve)
3. *d* < 48 μ*m* (*d* = width of the Gaussian-curve fit)
4. |*c*| < 24 μ*m*.

The points at the sperm head can be excluded from correction, if set by the user (Table 2K), which can improve the tracking of sperm with a head shape that does not fit well to a Gaussian curve. Next, outliers from the reconstructed flagellar track are removed: for each triplet of points (*p_i-1_*, *P_i_*, *p_i+1_*), *p_i_* is removed from *P* if 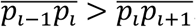. Afterwards, the flagellar track is smoothed by a custom-written algorithm: for a given point *p_i_* along the track, a distinct number (Table 2O) of upstream and downstream points from *p_i_* are included for smoothing. At the start and the end of the point list, less points are included. Pairs of the included points were generated in all possible combinations. For each pair, a line was constructed between the paired points and the point *p_i_* was projected to that line, resulting in the projected point *p_i_*. The median distance *p_i_* − *p_i_*’ of all pairs was determined and the flagellar track point *p_i_* was replaced with the projected point *p_i_*’ belonging to the pair whose projection distance was nearest to the median distance. Next, arc length positions *l_pi_*. for each flagellar point *p_i_* are calculated as: *l_pi_* = 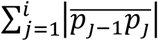. After smoothing, outliers – if still existing – are removed again and normal and tangential vectors are calculated as described. Again normal lines are constructed and the Gaussian-curve fits on the normal line are recalculated. The points with fits that do not match above described criteria can be removed, if set (Table 2H). For each point *p_i_* the width parameter of the fitted Gaussian curve serves as a relative z position of the flagellum (Figure 2G). The relative z position values are smoothed along the flagellum analogously to the described method for the smoothing in xy, except for that the points considered for smoothing upstream and downstream are restricted in the distance along the arc length to the smoothed point *p_i_* (Table 2N). Additionally, the Gaussian curve height is retrieved and saved as an intensity value for each flagellar point (Table 2H).

In the next step, all flagellar positions are translated into a new coordinate whose x axis is defined by the vector 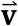 from the first point to the point at an indicated arc length position (Table 2P). The x and y positions of all flagellar points in the new coordinate system are plotted as a function of the arc length in kymographs (Figure 2D). Additionally, the angle between the x axis of the image and 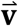 (angle θ) is output (Figure 2B). Further parameters are determined as described in the results part, e.g. the curvature angle (Figure 2F) is determined by the angle between the tangential vector at the given point and the tangential vector of the point in a defined arc length distance (Table 2Q) before the given point. The settings parameter “head rotation matrix radius” (Table 2T) determines the distance orthogonal to the sperm head that is included in calculating head torsion (Figure 2C). The FFT algorithm can not only be applied to the entire time sequence of a parameter but also for defined time steps of the image sequence; when the FFT parameter described in Table 2R is lower than the number of frames in the analysed time sequence, a window of the defined number of frames is slid along the time sequence and all FFT results are determined and output for each window separately. The sperm head can be excluded from the FFT analysis if it is not moving (e.g. in thethered sperm) (Table 2S).

### Software

Image processing and analysis were performed in ImageJ (U.S. National Institutes of Health, Bethesda, Maryland, USA). Plots and Figures were generated using GraphPad Prism (GraphPad Software, Inc., La Jolla, CA, USA) and Adobe Illustrator CS5 (Adobe Systems, Inc., San Jose, California, USA). The Java-based software was developed and compiled using Eclipse Mars.2 (IDE for Java Developers, Eclipse Foundation, Inc., Ottawa, Ontario, Canada).

### Software availability

*SpermQ* will be made accessible upon request. *SpermQ Evaluator* is publicly available (https://github.com/IIIImaging/SpermQ Evaluator).

## Acknowledgement

The author’s thank Dr. Luis Alvarez for critical discussions and reading of the manuscript. This work was supported by the German Research Foundation (DFG) in the priority program SPP 1726 “Microswimmers” (DW, JFJ) and by the Boehringer Ingelheim Fonds (JNH). DW is a member of the Excellence Cluster ImmunoSensation, Bonn.

## Author Contribution

JNH, JFJ, and DW designed the study. JNH developed software, prepared figures, analysed data, and drafted the manuscript. SR contributed to development of the *SpermQ Evaluator* tool. All authors wrote and edited the manuscript.

## Author Information

The authors declare no competing financial interests.

